# The Salas y Gomez and Nazca Ridges EBSA support a highly functional diversity of seabirds

**DOI:** 10.64898/2026.06.06.730628

**Authors:** Pamela Núñez, Guillermo Luna-Jorquera

## Abstract

The Salas y Gómez and Nazca Ridges (SGNRs) in the Southeast Pacific, recognized as an Ecologically or Biologically Significant Area (EBSA), host unique marine ecosystems with one of the highest rates of endemism on the planet. This study provides the first comprehensive trait-based assessment of seabird functional diversity in this globally significant region, focusing on their ecological contributions as top predators. Using at-sea abundance data from 11 oceanographic surveys (2014–2017) across 3,500 km of transects, we recorded 36 seabird species (8,179 individuals). We analysed functional diversity through ten foraging-related traits, including diet, foraging strata, and morphology. Multidimensional trait analyses revealed a seabird assemblage characterised by low functional richness (FRic = 0.0587), moderate-to-low evenness (FEve = 0.3649), and high divergence (FDiv = 0.6609), with non-random patterns confirmed by null models. Nesting (17 species) and non-nesting (19 species) groups showed distinct functional structures, with nesting seabirds exhibiting higher functional divergence and non-nesting seabirds greater functional evenness, though with 61% trait-space overlap. Low functional redundancy suggests that the loss of seabird species would likely translate into the loss of unique functional roles, potentially compromising ecosystem processes such as cross-system nutrient subsidies. With 73% of the SGNRs beyond national jurisdiction, seabirds face threats from unregulated fishing, plastic pollution, and seabed mining. These findings underscore the urgent need for conservation strategies under the High Seas Treaty (BBNJ treaty) to protect not only species richness but also functional roles, ensuring ecosystem resilience in this biodiversity hotspot of over 110 seamounts.

## Introduction

The Salas y Gómez and Nazca Ridges (SGNRs) in the Southeast Pacific stand as a globally recognized Ecologically or Biologically Significant Area (EBSA), distinguished by one of the highest rates of marine endemism on seamounts worldwide^1^. An Ecologically or Biologically Significant Marine Area (EBSA) is a part of the ocean identified as having special ecological or biological importance, based on scientific criteria, to support conservation and sustainable management efforts^1^. Further studies supporting the SGNR EBSA designation^2,3^ acknowledge seabirds as part of biodiversity, yet their treatment remains taxonomic rather than functional. This region hosts unique deep-sea ecosystems associated with its extensive seamount ridges and complex oceanographic features, including submarine canyons and benthic habitats supporting rich invertebrate communities. It also constitutes critical pelagic habitats serving as migration corridors for species like Blue whale (*Balaenoptera musculus*) while supporting over 80 threatened or endangered taxa, including sea turtles and seabirds, and serving as a nursery for economically important species such as Chilean jack mackerel (*Trachurus murphyi*) and swordfish (*Xiphias* gladius)^2^. Beyond their economic value, pelagic forage species such as jack mackerel occupy a low– to mid-trophic-level position in the regional food web, channelling energy from plankton and smaller prey to higher trophic levels and thereby sustaining top predators, including seabirds. In addition to the ecological role, the region’s status as a natural and cultural heritage site, tied to centuries of Polynesian seafaring, underscores its broader significance^2^. Due to the region’s ecological, biological and cultural value, Chile established four Marine Protected Areas around the islands associated with the seamount ridges of Nazca and Salas & Gómez, while Peru implemented a marine area to safeguard the seabed (Figure 1)^4,5^. However, 73% of the region lies beyond national jurisdictions, where current governance is insufficient to address global threats such as climate change and plastic pollution, as well as regional pressures including overfishing and potential seabed mining^2^. These challenges are driving international efforts, such as the recently adopted High Seas Treaty –formally, the Agreement on Biodiversity Beyond National Jurisdiction (BBNJ Treaty)– to protect biodiversity in areas outside national waters^2^.

**Figure 1.**
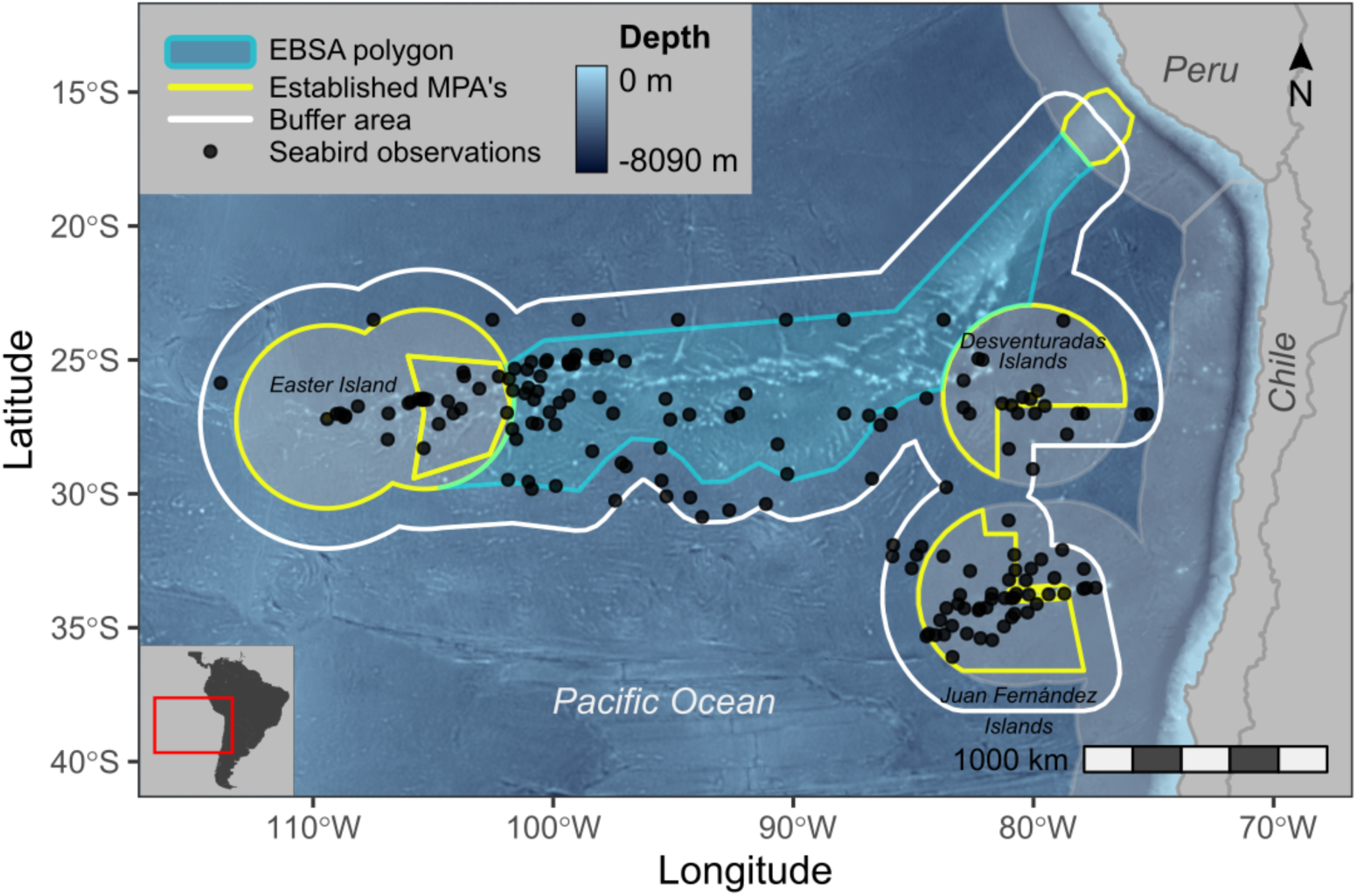
The map showing the study area (white line) includes the Salas & Gómez and Nazca ridges EBSA and diverse marine protected areas around Chilean oceanic islands (see text for details). It shows the Peruvian and Chilean Exclusive Economic Zones and the seafloor bathymetry. Black dots show the position of the daily surveys to determine seabird richness and abundance of seabirds at sea (see the text for details). The map was plotted using the R package ggplot2^46^.

Seabirds are central to the ecological dynamics of the SGNRs, acting as top predators and indicators of marine health across the Southeast Pacific^6^. Recent studies reveal a distinct biogeographic pattern, with significant species composition and diversity variation among seabird assemblages nesting on coastal and oceanic islands. Species richness declines from Chilean coastal islands to distant Polynesian islands, shaped by gradients in island physiography and oceanographic factors like primary productivity^7^. This pattern reflects a structured gradient of species replacement (β-turnover) across systems like the Galápagos and Chilean islands rather than nested subsets, indicating specific adaptations to environmental filters^7^. Within this context, the SGNRs provide essential nesting habitats on islands and extensive foraging grounds at sea, supporting diverse seabird communities^6,8^, including long-distance migrants that travel thousands of kilometres between hemispheres during breeding and non-breeding phases^9,10^.

As top predators, seabirds exhibit diverse ecological strategies to exploit various foraging niches, significantly contributing to ecosystem processes like nutrient cycling and energy transfer between terrestrial and marine environments^11,13^. Their functional diversity, encompassing a range of traits related to resource use, underpins ecosystem resilience and stability^14,16^. Research in the Humboldt Current System shows that oceanographic gradients foster niche differentiation, promoting assemblages with limited functional overlap^17,18^. These studies further indicate that multi-species feeding flocks are shaped by intrinsic traits like taxonomic affinity and foraging guilds rather than just food availability, highlighting the role of behavioural differences in shaping the structure of these assemblages^17,18^. Such patterns suggest that each species plays a distinct role, enhancing ecosystem function but increasing vulnerability to loss, especially under threats like overfishing and climate change that may disrupt foraging and breeding success.

Despite their likely importance in the SGNRs, the specific contributions of seabirds to ecosystem functioning remain underexplored. The SGNRs are an extremely isolated system, bounded by major oceanographic features (*i.e.*, the Atacama Trench and the Humboldt Current System) and an oxygen minimum zone, which generate high endemism, a strongly filtered species pool, and pronounced gradients in productivity, temperature, and oxygen^2^. To explore the ecological roles of seabirds in the SGNRs, we focus on functional diversity through selected traits that primarily reflect foraging ecology (*e.g.*, diet, foraging stratum, flight-related morphology) while also capturing broader functional variation related to movement capacity, energy requirements, and general ecological strategies^19,21^.

We hypothesized (***i***) that the functional diversity of the seabird assemblage in the SGNRs is defined by low functional richness and an uneven distribution of ecological roles, resulting from a constrained range of functional traits and pronounced ecological structuring, and (***ii***) that functional diversity differs significantly between nesting and non-nesting seabird groups in the SGNRs, with non-nesting seabirds displaying greater functional richness and evenness, and nesting seabirds exhibiting higher functional divergence. This pattern is expected because nesting species are subject to local constraints, *e.g.*, breeding site fidelity and specific resource use, which select for particular morphological and lifestyle trait combinations (*i.e.*, body mass, wing, tarsus, beak and tail lengths, and foraging lifestyle). These constraints arise, for example, because breeding seabirds, as central place foragers, must incubate and feed their chicks at the colony, restricting their foraging range. As a result, niche separation may be especially important when multiple species need to exploit resources within the same area^22,23^. In contrast, non-nesting species represent a more heterogeneous set of ecological strategies and morphologies, reflecting diverse evolutionary histories and broader trait variability within the regional assemblage.

In this study, we defined nesting species as seabirds that breed within the SGNRs, predominantly including taxa such as frigatebirds, boobies, terns, and petrels that maintain established breeding colonies in the study area. In contrast, non-nesting species comprised seabirds present in the study area, but which do not breed locally, typically including various albatrosses, large petrels, and other migratory or transient taxa observed at sea whose breeding sites lie outside the SGNRs. In total, our assemblage included 17 nesting and 19 non-nesting species. Therefore, our main objective was to analyse the functional diversity of seabird assemblages by constructing a multidimensional functional trait space for seabirds of the SGNRs and comparing diversity metrics between nesting and non-nesting groups, aiming to clarify their roles in marine ecological processes and support conservation strategies for this globally significant region.

## Methods

### Study Area

We assessed seabird functional diversity within the EBSA of the SGNRs in the Southeast Pacific Ocean (Figure 1). This region comprises two contiguous volcanic seamount chains extending approximately 2,900 km from the westernmost end of the Salas y Gómez Ridge to the northeastern edge of the Nazca Ridge near the Peru-Chile Trench^24,25^. The SGNRs include over 110 seamounts located in areas beyond national jurisdiction (ABNJ), with summit depths ranging from 1,000 to 3,400 m^26^. Oceanographic conditions in the region are shaped by the South Pacific Subtropical Gyre, which includes nutrient pulses from Taylor column circulations, promoting local upwelling and enhanced productivity^27,28^.

To define our study area, we combined the SGNRs EBSA polygon^29^ with established Marine Protected Areas (MPAs) within the Exclusive Economic Zones of Chile^4^ and Peru^5^. These MPAs include: (***i***) the Rapa Nui Multipurpose Coastal Marine Area and Motu Motiro Hiva Marine Park (Rapa Nui and Salas y Gómez islands); (***ii***) the Nazca-Desventuradas Marine Park (Desventuradas Islands); (***iii***) the Mar de Juan Fernández and Montes Submarinos Crusoe y Selkirk Marine Park (Juan Fernández islands); and (***iv***) the Reserva Nacional Dorsal de Nazca (Peru). We applied a 156 km buffer around the merged EBSA-MPA polygons to encompass the foraging ranges of key seabird species known to nest on oceanic islands and forage widely across this region^6,30^. This buffer width was determined as the minimum distance required to fully enclose all EBSA and MPA polygons within a single spatial unit, ensuring a coherent ecological area for analysis. The resulting polygon was used to extract seabird transects and define the geographic scope of all functional diversity analyses. This spatial extent was selected to capture both the core breeding areas and foraging ranges of seabirds using the SGNR region, enabling the analysis of trait diversity across a meaningful ecological scale.

### Seabird Occurrence and Abundance Data Collection

Data on seabird occurrence and abundance at sea were obtained from the Seabirds at Sea Monitoring Program (SASMP; see Serratosa *et al*.^6^ for full methodology). Observations were conducted during 11 oceanographic surveys between 2014 and 2017, and although not every month was sampled in each year, the combined surveys encompassed all months, thereby completing a full annual cycle along an oceanographic transect of approximately 3,500 km from the Chilean continental margin to Rapa Nui Island, following different but partially overlapping routes among cruises. Seabirds were recorded using standardized strip transect methods. An experienced observer, positioned on the vessel’s flying bridge, continuously recorded all seabirds within a 300-meter strip on one side of the ship (a 90° arc from bow to beam). Data were collected in continuous 10-minute intervals. For analysis, all 10-minute intervals recorded on the same calendar day were aggregated into a single daily transect, which served as the sampling unit. To account for variation in sampling effort across daily transects, seabird counts were standardized by the area examined during a day to obtain density estimates (individuals·km⁻^2^), which we refer to as abundance throughout this study. Transects with no bird observations or with records only identified to genus level (*e.g.*, *Pterodroma* sp.) were excluded. In total, 167 daily transects were recorded within the study area, resulting in observations of 36 seabird species and 8,179 individuals. A seabird species was classified as nesting or non-nesting based on confirmed breeding activity on nearby oceanic islands^7,30,31^.

Species richness and Chao1 diversity index were calculated for each daily transect using the *vegan* R package^32^. Sampling completeness was assessed by generating a species accumulation curve with the *specaccum()* function and estimating asymptotic richness using the Chao1 estimator via the *ChaoSpecies()* function in the *SpadeR* package^33^. This method estimates total richness based on the number of rare species, defined as those observed only once or twice. Rare species were categorized as those with ≤10 individuals across all transects, a threshold chosen to conservatively include species with low but non-negligible representation, reflecting the variability in transect effort and ensuring robust estimation of sampling completeness. Confidence intervals (95%) were calculated separately for rare and abundant species groups to evaluate the completeness of our sampling.

## Functional Diversity

### Functional Traits

The traits we used here were selected because they reflect species’ life history and are established proxies for ecological function in seabirds^21,30^. The morphological measurements represent evolutionary adaptions associated with prey capture, foraging mode, and dispersal capacity^34^, providing a quantitative basis for distinguishing functional roles among species. We selected five standardized continuous morphological traits: beak length (culmen) (mm), tarsus length (mm), wing length (mm), tail length (mm), and body mass (g); and then performed a Principal Component Analysis (PCA) to reduce redundancy among these correlated traits. The first two principal components explained 97% of the total variance and were retained as composite axes representing overall body size and shape variation. These axes were used as continuous inputs in the functional trait space.

In addition to continuous traits, we included ecological traits that were either categorical or fuzzy-coded (Table S1). Categorical traits included primary lifestyle (aerial, aquatic, or generalist) coded as nominal as these categories reflect alternative ecological strategies without a functional hierarchy, and diet type (invertivore, omnivore, or piscivore/scavenger), coded as ordinal following a general trophic gradient: Invertebrate < Omnivore < Vertebrate/Fish Scavenger. The invertivore category encompasses species that consume both planktonic organisms and other invertebrates such as cephalopods, as insufficient detailed dietary data exist for all species to distinguish between planktivorous and nekton-feeding strategies. Fuzzy-coded traits described the proportional association of each species with three foraging strata: below the surface, around the surface, and on the ground, summing to one per species. Trait values were extracted from Tobias *et al.*^34^ for morphology and lifestyle, and from Wilman *et al.*^35^ for foraging strata and diet. Primary lifestyle definitions follow Tobias *et al.*^34^, which are based on predominant habitat and activity patterns during the annual cycle. During southward migration, the Red Phalarope exhibits pelagic habits and feeds primarily on marine invertebrates^36^; therefore, we classified it as an aquatic invertivore within the study area.

### Functional distances

To compute pairwise functional distances among species, we prepared two input tables following the requirements of the mFD package^37^: (***i***) a species-by-trait matrix that included the two PCA-derived axes (PC1-size variation, PC2-shape variation), the categorical traits (Diet.5Cat and Primary Lifestyle), and the fuzzy-coded foraging strata, and (***ii***) a trait type metadata table that specified the nature of each trait as quantitative (Q), ordinal (O), nominal (N), or fuzzy (F) as appropriate, and grouped the fuzzy traits under a shared set (“Foraging”). Then we quantified the functional distances using the Gower dissimilarity matrix with the *funct.dist()* function from the package mFD. Prior to distance calculation, the quantitative traits were centered and scaled (z-scores) to ensure comparability.

### PCoA and Space Quality

We constructed a multidimensional functional space by applying a Principal Coordinates Analysis (PcoA) to the species-by-species Gower matrix previously obtained. We retained the 14 axes with positive eigenvalues for all functional diversity metrics and quality evaluations. Since Gower distances are not strictly Euclidean, the relative eigenvalues from the PcoA summed to more than one. To report variance explained by the main axes, we normalized the eigenvalues so their sum equalled one. This normalization was used only for interpretation and did not affect the space construction. The quality of spaces with increasing dimensionality (2 to 14 axes) was evaluated using the *quality.fspaces()* function, with mean absolute deviation (MAD) as the performance criterion. Subsequent analyses were based on the first three PcoA axes, which were identified as providing optimal space quality as they offered an optimal balance between dimensionality and space quality. We assessed the contribution of individual traits to the principal axes of the functional space using the *traits.faxes.cor()* function. Trait–axis associations were quantified using either linear models (R²) for continuous and fuzzy traits, or Kruskal–Wallis tests (η²) for categorical traits. Traits with R² or η² values greater than 0.5 were considered strong contributors, while those less than 0.2 were considered weakly associated. Significance (p < 0.05) was assessed via permutation procedures implemented within the function. To visualize the functional space, we used *the funct.space.plot()* function from the mFD package to project species coordinates across all pairwise combinations of the first three PcoA axes, facilitating ecological interpretation of the trait space.

### Functional Diversity Indices

We quantified functional diversity at two levels within the SGNRs seabird assemblage: (***i***) the entire seabird assemblage and, (***ii***) behavioural groups defined by nesting status. To describe overall functional structure, we first aggregated species-specific densities across all transects, producing a single abundance value per species and a community matrix (species × abundance) for the full assemblage. Functional diversity was then computed for the full assemblage of 36 species, and separately for the 17 nesting and 19 non-nesting species.

We used four complementary indices to capture different facets of functional structure in trait space. Functional Richness (FRic) quantifies the volume of trait space occupied by the community. Functional Evenness (FEve) measures the regularity of species distributions, while Functional Divergence (FDiv) reflects the dominance of species with extreme trait values^38,39^. Rao’s quadratic entropy (RaoQ) integrates pairwise trait dissimilarity weighted by species abundance^40,41^. FRic, FEve, and FDiv were calculated using the *alpha.fd.multidim()* function from the mFD R package^37^, based on species coordinates from the three PcoA axes selected according to space quality (see PcoA and Space Quality). RaoQ was computed using the *rao.diversity()* function from SYNCSA^42^, based on the Gower matrix combining continuous, categorical, and fuzzy traits. To visualize the functional indices we used the *alpha.multidim.plot()* function from mFD^37^.

To quantify functional similarity between nesting and non-nesting species, we estimated trait-space overlap using a 3D kernel density estimation^43^ and the Bhattacharyya Affinity index^44^. This index accounts for both the extent and intensity of shared trait-space use, with values above 0.5 indicating substantial functional overlap.

Finally, we used null models to test (***i***) whether observed trait distributions in the complete seabird assemblage were more or less structured than expected by chance, and (***ii***) whether differences in functional diversity exist between nesting and non-nesting seabirds. For both tests, we ran 999 iterations, fixing species identities and abundances while randomizing species. For each iteration, we recalculated a Gower distance matrix, constructed a functional space using the first three PcoA axes, and computed Functional Richness (FRic), Evenness (FEve), and Divergence (FDiv) with the mFD package^37^. Rao’s quadratic entropy (RaoQ) was calculated using the *rao.diversity()* function from the SYNCSA package^42^, based on randomized Gower distances. Two-tailed p-values and standardized effect sizes (SES) were used to assess deviations from random expectations. When significant differences were found between groups for a given functional diversity metric, we quantified each species’ contribution by recalculating the index after removing each species in turn (“leave-one-out” approach). We then expressed the absolute change as a percentage of the original group value, which allowed us to rank species by their relative influence on the metric within each group.

We performed all analyses in the R software^45^ and its associated packages. All figures were plotted using the R package ggplot2^46^.

## Results

### Richness, diversity and abundance

The species accumulation curve approached an asymptote after ∼150 sampling days, indicating that sampling effort was sufficient to capture the area’s seabird diversity (Supplementary Figure S1). Chao1 estimators predicted a total species richness of 37 (95% CI: 36–47), closely matching the observed richness of 36 species (Table 1), and confirming near-complete sampling coverage (∼95% for rare species). Species richness and Chao1 diversity per transect showed spatial variability (Figure 2a and b). Richness ranged from 1 to 11 species per transect (mean = 3.5, median = 3), while Chao1 diversity varied from 0 to 2.08 (mean = 0.75, median = 0.69). Spatial patterns highlighted areas of elevated species richness and Chao1 diversity within the EEZs around the islands, although some higher values were observed within the SGNR.

**Figure 2.**
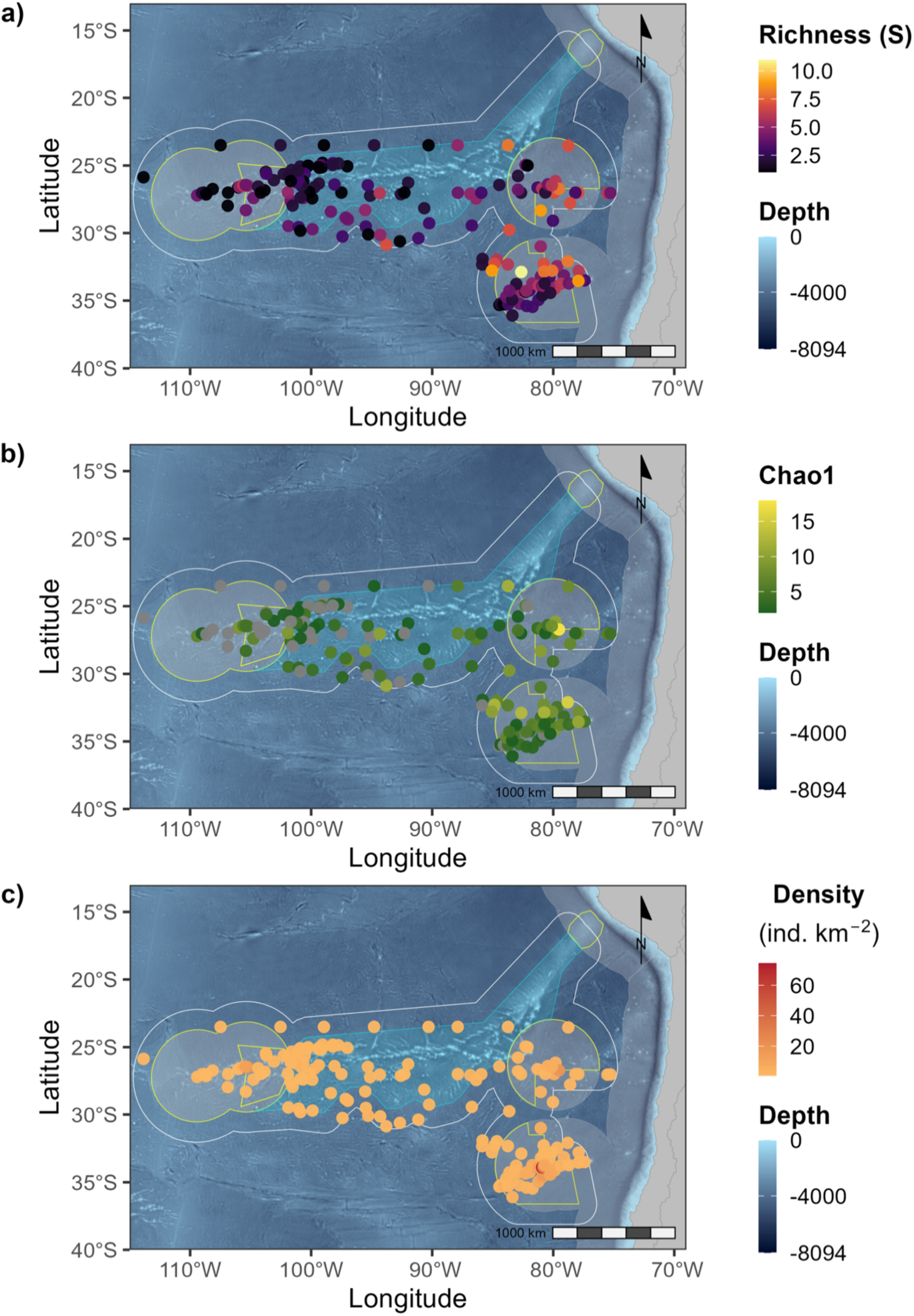
Spatial distribution of seabird: **a)** species richness (S), **b)** Chao1 diversity, and **c)** abundance (km^-2^) across transects in the study area. The maps were plotted using the R package ggplot2^46^.

**Table 1.**
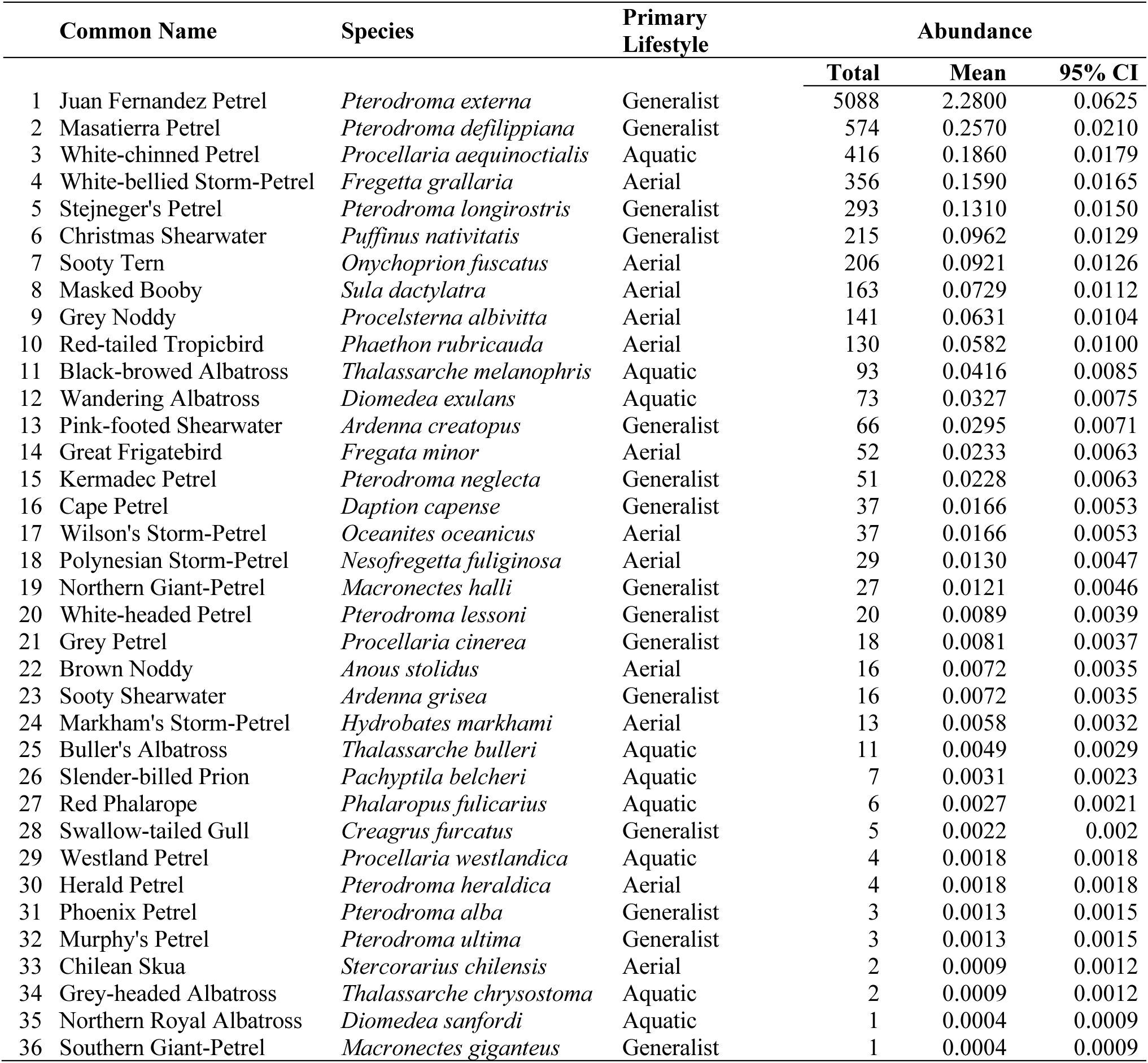
Seabird species registered in the Salas & Gómez and Nazca Ridges. ordered by abundance. Total is the number of birds recorded across 167 daily transects. The mean is the number of birds per total area surveyed (bird·km^-2^); and “95% CI” is the Poisson confidence interval for count data. For details on the primary lifestyle classification see Materials and Methods.

|  | Common Name | Species | Primary Lifestyle | Abundance |  |  |
| --- | --- | --- | --- | --- | --- | --- |
|  |  |  |  | Total | Mean | 95% CI |
| 1 | Juan Fernandez Petrel | *Pterodroma externa* | Generalist | 5088 | 2.2800 | 0.0625 |
| 2 | Masatierra Petrel | *Pterodroma defilippiana* | Generalist | 574 | 0.2570 | 0.0210 |
| 3 | White-chinned Petrel | *Procellaria aequinoctialis* | Aquatic | 416 | 0.1860 | 0.0179 |
| 4 | White-bellied Storm-Petrel | *Fregetta grallaria* | Aerial | 356 | 0.1590 | 0.0165 |
| 5 | Stejneger's Petrel | *Pterodroma longirostris* | Generalist | 293 | 0.1310 | 0.0150 |
| 6 | Christmas Shearwater | *Puffinus nativitatis* | Generalist | 215 | 0.0962 | 0.0129 |
| 7 | Sooty Tern | *Onychoprion fuscatus* | Aerial | 206 | 0.0921 | 0.0126 |
| 8 | Masked Booby | *Sula dactylatra* | Aerial | 163 | 0.0729 | 0.0112 |
| 9 | Grey Noddy | *Procelsterna albivitta* | Aerial | 141 | 0.0631 | 0.0104 |
| 10 | Red-tailed Tropicbird | *Phaethon rubricauda* | Aerial | 130 | 0.0582 | 0.0100 |
| 11 | Black-browed Albatross | *Thalassarche melanophris* | Aquatic | 93 | 0.0416 | 0.0085 |
| 12 | Wandering Albatross | *Diomedea exulans* | Aquatic | 73 | 0.0327 | 0.0075 |
| 13 | Pink-footed Shearwater | *Ardenna creatopus* | Generalist | 66 | 0.0295 | 0.0071 |
| 14 | Great Frigatebird | *Fregata minor* | Aerial | 52 | 0.0233 | 0.0063 |
| 15 | Kermadec Petrel | *Pterodroma neglecta* | Generalist | 51 | 0.0228 | 0.0063 |
| 16 | Cape Petrel | *Daption capense* | Generalist | 37 | 0.0166 | 0.0053 |
| 17 | Wilson's Storm-Petrel | *Oceanites oceanicus* | Aerial | 37 | 0.0166 | 0.0053 |
| 18 | Polynesian Storm-Petrel | *Nesofregetta fuliginosa* | Aerial | 29 | 0.0130 | 0.0047 |
| 19 | Northern Giant-Petrel | *Macronectes halli* | Generalist | 27 | 0.0121 | 0.0046 |
| 20 | White-headed Petrel | *Pterodroma lessoni* | Generalist | 20 | 0.0089 | 0.0039 |
| 21 | Grey Petrel | *Procellaria cinerea* | Generalist | 18 | 0.0081 | 0.0037 |
| 22 | Brown Noddy | *Anous stolidus* | Aerial | 16 | 0.0072 | 0.0035 |
| 23 | Sooty Shearwater | *Ardenna grisea* | Generalist | 16 | 0.0072 | 0.0035 |
| 24 | Markham's Storm-Petrel | *Hydrobates markhami* | Aerial | 13 | 0.0058 | 0.0032 |
| 25 | Buller's Albatross | *Thalassarche bulleri* | Aquatic | 11 | 0.0049 | 0.0029 |
| 26 | Slender-billed Prion | *Pachyptila belcheri* | Aquatic | 7 | 0.0031 | 0.0023 |
| 27 | Red Phalarope | *Phalaropus fulicarius* | Aquatic | 6 | 0.0027 | 0.0021 |
| 28 | Swallow-tailed Gull | *Creagrus furcatus* | Generalist | 5 | 0.0022 | 0.002 |
| 29 | Westland Petrel | *Procellaria westlandica* | Aquatic | 4 | 0.0018 | 0.0018 |
| 30 | Herald Petrel | *Pterodroma heraldica* | Aerial | 4 | 0.0018 | 0.0018 |
| 31 | Phoenix Petrel | *Pterodroma alba* | Generalist | 3 | 0.0013 | 0.0015 |
| 32 | Murphy's Petrel | *Pterodroma ultima* | Generalist | 3 | 0.0013 | 0.0015 |
| 33 | Chilean Skua | *Stercorarius chilensis* | Aerial | 2 | 0.0009 | 0.0012 |
| 34 | Grey-headed Albatross | *Thalassarche chrysostoma* | Aquatic | 2 | 0.0009 | 0.0012 |
| 35 | Northern Royal Albatross | *Diomedea sanfordi* | Aquatic | 1 | 0.0004 | 0.0009 |
| 36 | Southern Giant-Petrel | *Macronectes giganteus* | Generalist | 1 | 0.0004 | 0.0009 |

Seabird abundance varied widely with mean bird density per day ranging from 0.004 to 2.28 bird·km⁻² (Figure 2c). The most abundant species were the Juan Fernández Petrel, Masatierra Petrel, and White-chinned Petrel. The least abundant species were the Northern Royal Albatross and Southern Giant Petrel (Table 1).

### Functional space

We constructed a three-dimensional functional space based on a principal coordinate analysis (PcoA) of Gower distances calculated from seabird trait data, including categorical, continuous and fuzzy variables. The 3D solution minimized mean absolute deviation (MAD = 0.033), providing an optimal representation of trait-based distances (Table 2; Supplementary Figure S2). PC1 (40.2% of normalized variance) was primarily associated with the categorical diet trait (η² = 76.9%) and, from the three foraging strata, was most strongly influenced by the proportion of foraging done around the water surface (R² = 51.1%). PC2 (30.3%) reflected differences in primary lifestyle (η² = 86.8%), with minor contributions from diet (η² = 17.4%). PC3 (14.0%) was strongly influenced by body size (PC1 morph; R² = 82.4%) and lifestyle (η² = 40.0%) (Table 2; Figure S3). The functional trait space of the entire seabird assemblage showed a relatively compact structure, with moderate expansion along PC1 (diet-related traits) and PC2 (lifestyle; Figure 3). Of the 36 species, 16 occupied the vertices of the convex hull, representing extreme trait combinations, while the remaining 20 species clustered within intermediate regions of the functional space.

**Figure 3.**
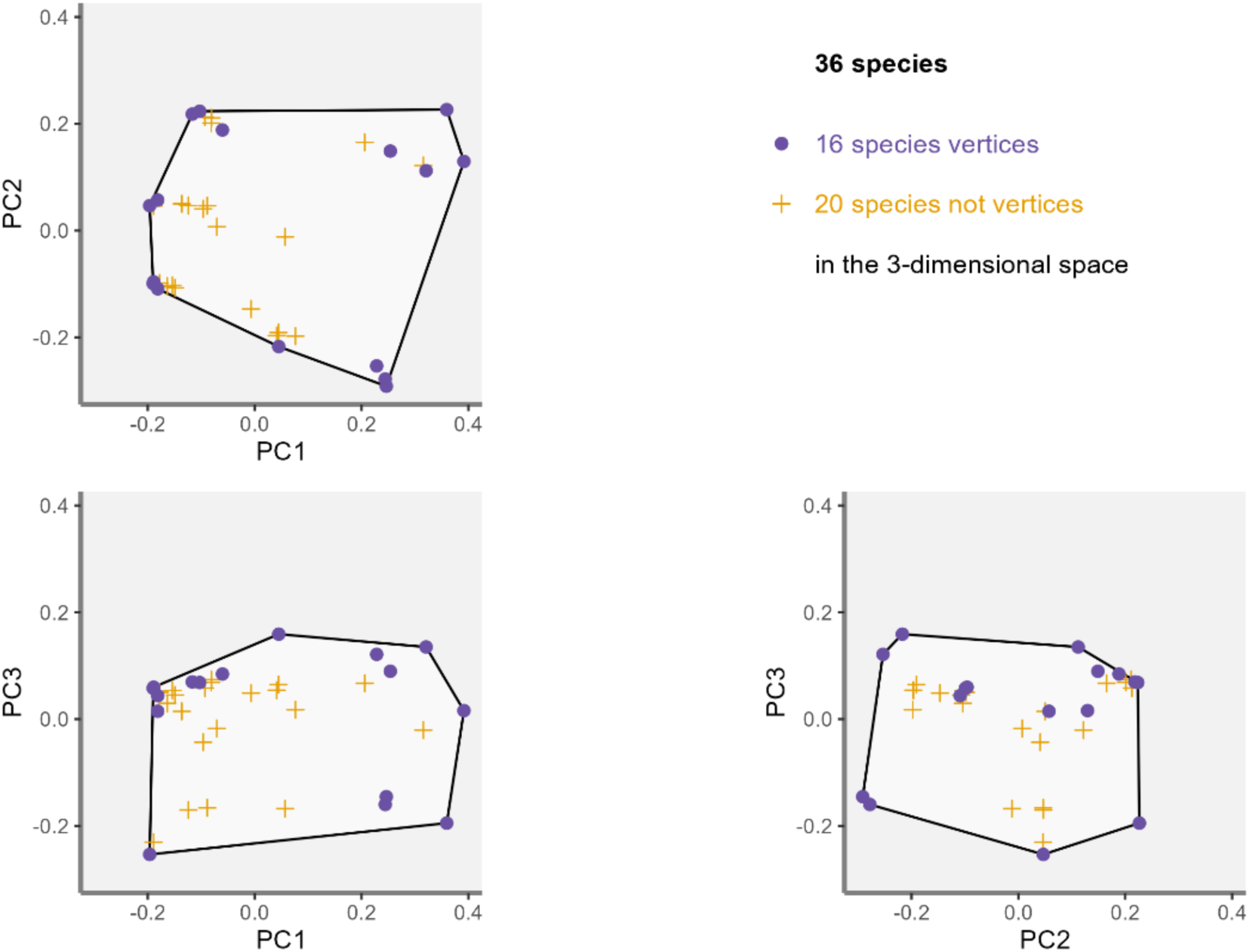
Functional trait space of the full seabird assemblage in the SGNRs EBSA projected onto the first three PCoA axes. Biplots show combinations of (PC1 vs PC2), (PC1 vs PC3), and (PC2 vs PC3). Black convex hulls delineate the boundaries of the functional space, with purple points indicating species at the vertices (extreme trait combinations) and yellow crosses representing remaining species.

**Table 2.**
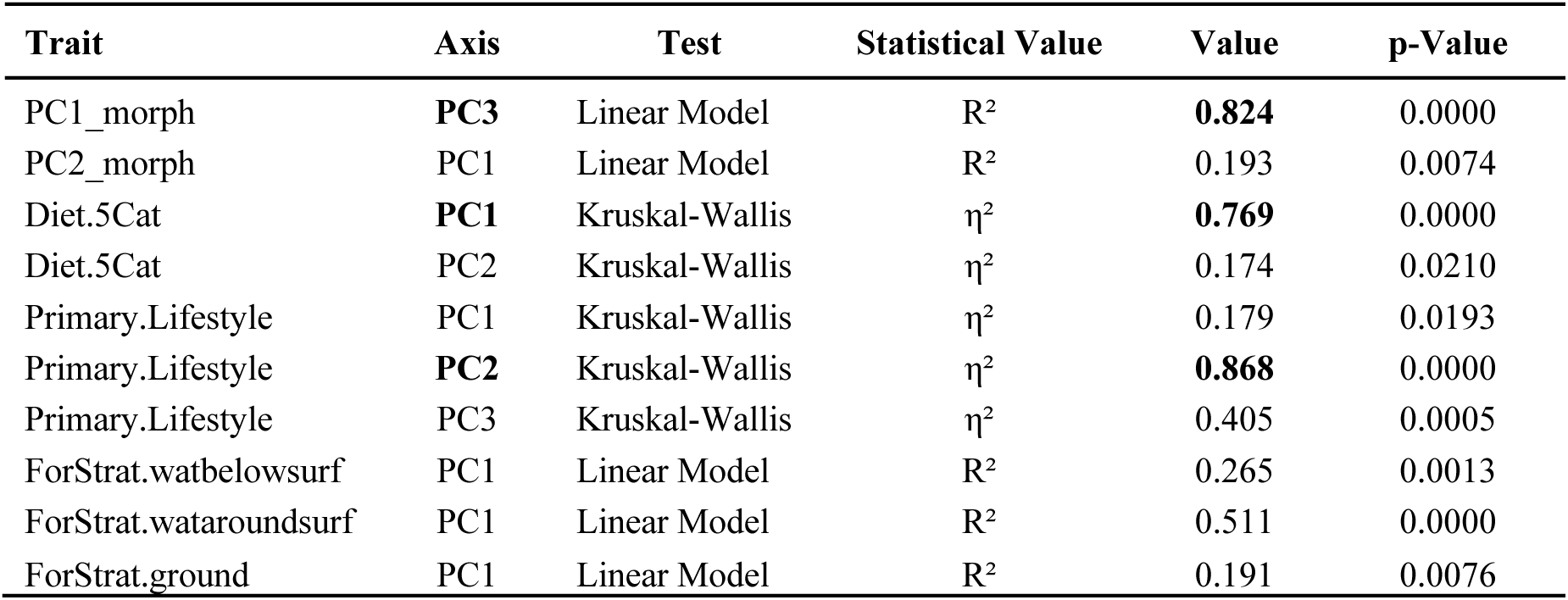
Relationships between PCoA axes and seabird ecological traits in the SGNRs EBSA. Statistical metrics include the coefficient of determination (R²) for linear models and effect size (η²) for Kruskal-Wallis tests. P-values indicate statistical significance (p < 0.05). Bold values highlight the strongest associations.

| Trait | Axis | Test | Statistical Value | Value | p-Value |
| --- | --- | --- | --- | --- | --- |
| PC1_morph | **PC3** | Linear Model | $R^2$ | **0.824** | 0.0000 |
| PC2_morph | PC1 | Linear Model | $R^2$ | 0.193 | 0.0074 |
| Diet.5Cat | **PC1** | Kruskal-Wallis | $\eta^2$ | **0.769** | 0.0000 |
| Diet.5Cat | PC2 | Kruskal-Wallis | $\eta^2$ | 0.174 | 0.0210 |
| Primary.Lifestyle | PC1 | Kruskal-Wallis | $\eta^2$ | 0.179 | 0.0193 |
| Primary.Lifestyle | **PC2** | Kruskal-Wallis | $\eta^2$ | **0.868** | 0.0000 |
| Primary.Lifestyle | PC3 | Kruskal-Wallis | $\eta^2$ | 0.405 | 0.0005 |
| ForStrat.watbelowsurf | PC1 | Linear Model | $R^2$ | 0.265 | 0.0013 |
| ForStrat.wataroundsurf | PC1 | Linear Model | $R^2$ | 0.511 | 0.0000 |
| ForStrat.ground | PC1 | Linear Model | $R^2$ | 0.191 | 0.0076 |

### Functional Diversity at the Assemblage Level

For the full assemblage, functional richness was relatively low (FRic = 0.0587), indicating broad occupation of niche space, whereas functional evenness was moderate (FEve = 0.3649), functional divergence was high (FDiv = 0.6609), and Rao’s quadratic entropy also indicated high functional dissimilarity (RaoQ = 0.7546). The functional trait space of the entire seabird assemblage showed a relatively compact structure, with moderate expansion along PC1 (diet-related traits) and PC2 (lifestyle; Figure 3). Null model results indicated that the observed value of FRic was significantly lower than expected under random assembly, and both FEve and FDiv were also outside the range expected by chance, reflecting a non-random functional structure in the community.

### Functional Diversity at the Level of Nesting and Non-nesting Seabirds

In the multidimensional trait space, nesting and non-nesting seabirds occupied largely overlapping regions but differed in how they filled that space (Figures 4 to 6). Projections of functional richness, evenness and divergence showed that both groups shared most of the global functional space, while a subset of species in each group extended their convex hulls towards non-overlapping margins. We observed that for each metric (except for RaoQ), the convex hulls of nesting and non-nesting seabirds overlap in all biplots by approximately 61%, as confirmed by the Bhattacharyya Affinity index (BA = 0.61). These spatial patterns were consistent with the functional diversity metrics (Figure 7); nesting species showed greater functional divergence, while non-nesting species exhibited higher functional richness and evenness. Direct comparison between nesting and non-nesting groups revealed that the observed difference in FRic was significantly lower than expected under the null model (SES >> 1.96), as was the difference for FEve (SES = –2.43). No significant differences between these groups were detected for FDiv (SES = 1.91) or RaoQ (SES = –0.16).

**Figure 4.**
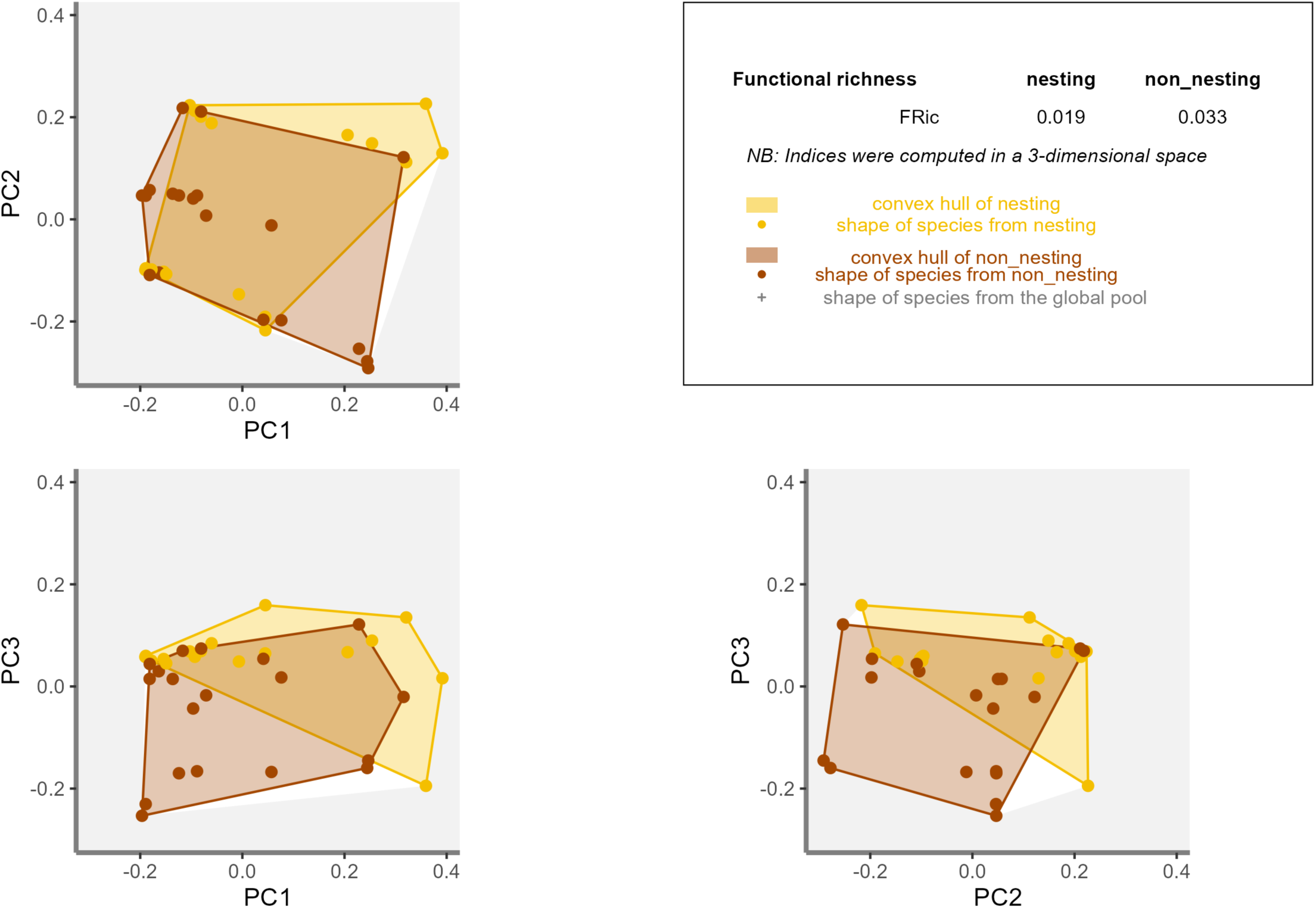
Functional richness (FRic) of nesting and non-nesting seabird groups projected onto three principal coordinate axes (PC1–PC2, PC1–PC3, and PC2–PC3). The white polygon represents the global functional space defined by all seabird species (species pool). Green and yellow points correspond to nesting and non-nesting species, respectively. Coloured surfaces display the convex hulls of each group, representing their occupied trait space. Despite substantial overlap between groups, several species contributed to marginal differentiation at the edges of the functional space (see Table 4).

**Figure 5.**
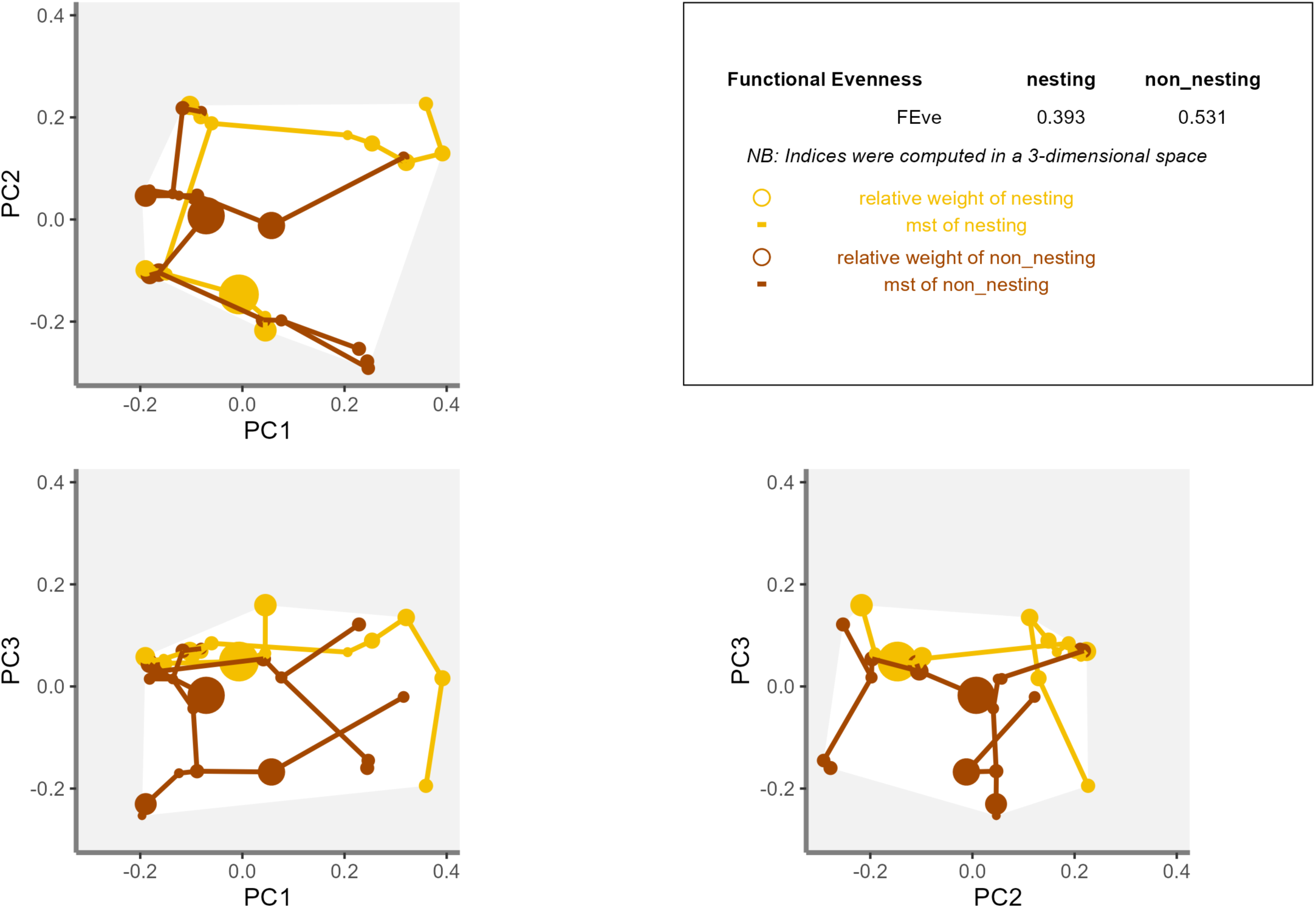
Functional evenness (FEve) of nesting (orange) and non-nesting (brown) seabird groups within the global functional space. FEve describes the regularity of species distribution within the occupied trait space. The figure shows species positions (point size proportional to species relative abundance within each group), the minimum spanning tree (MST) connecting species within each group, and the convex hull (white) representing functional richness (FRic). Higher FEve values indicate longer MST branches and a more even distribution of species across the functional space.

**Figure 6.**
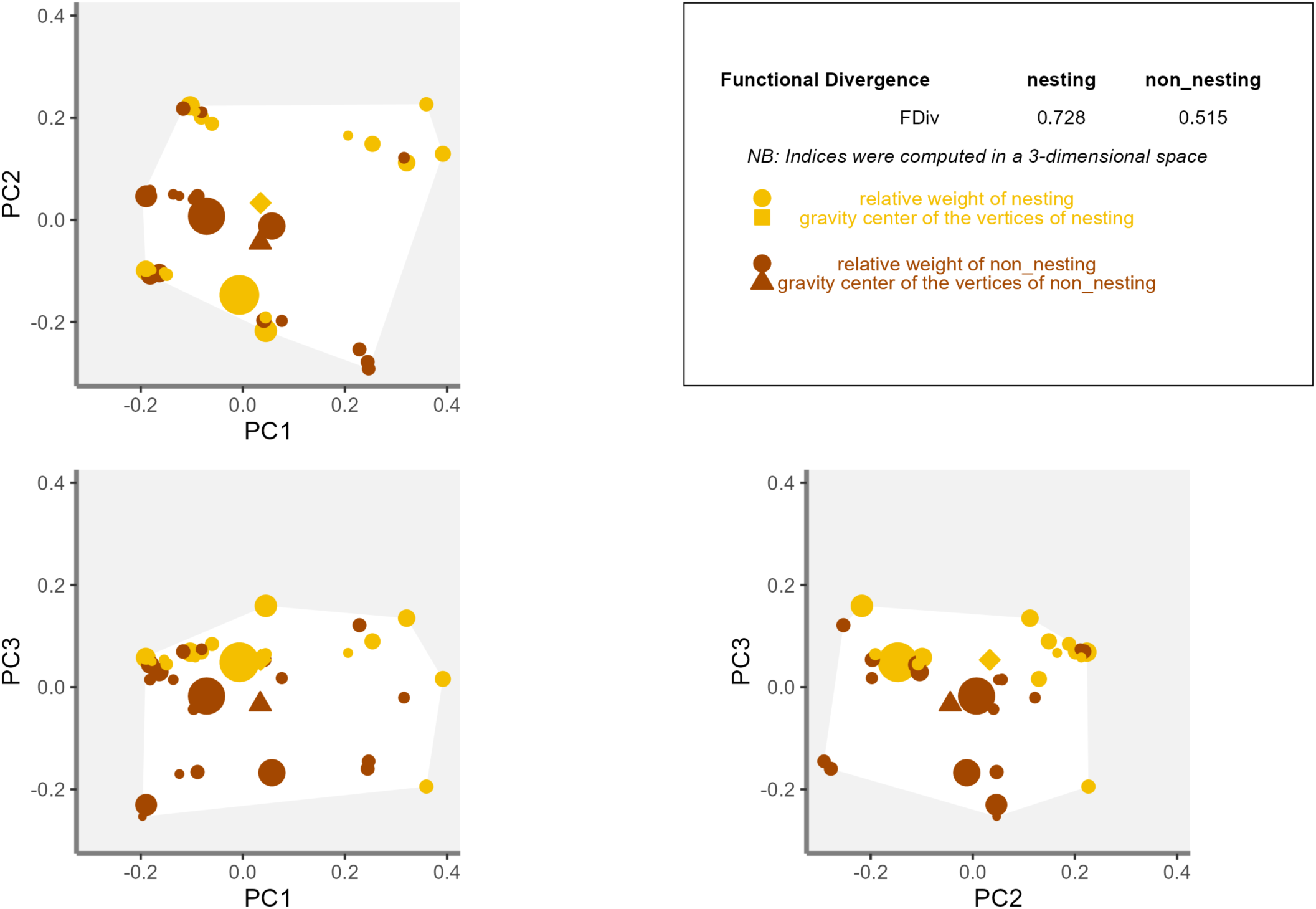
Functional divergence (FDiv) of nesting (orange) and non-nesting (brown) seabird groups within the global functional space. FDiv measures the extent to which species are distributed towards the edges of the functional space. The figure shows species positions (point size proportional to species relative abundance within each group), the centroid (gravity center) of species positions, and the convex hull (white) representing functional richness (FRic). Higher FDiv values indicate a greater contribution of marginal species located far from the centroid.

**Figure 7.**
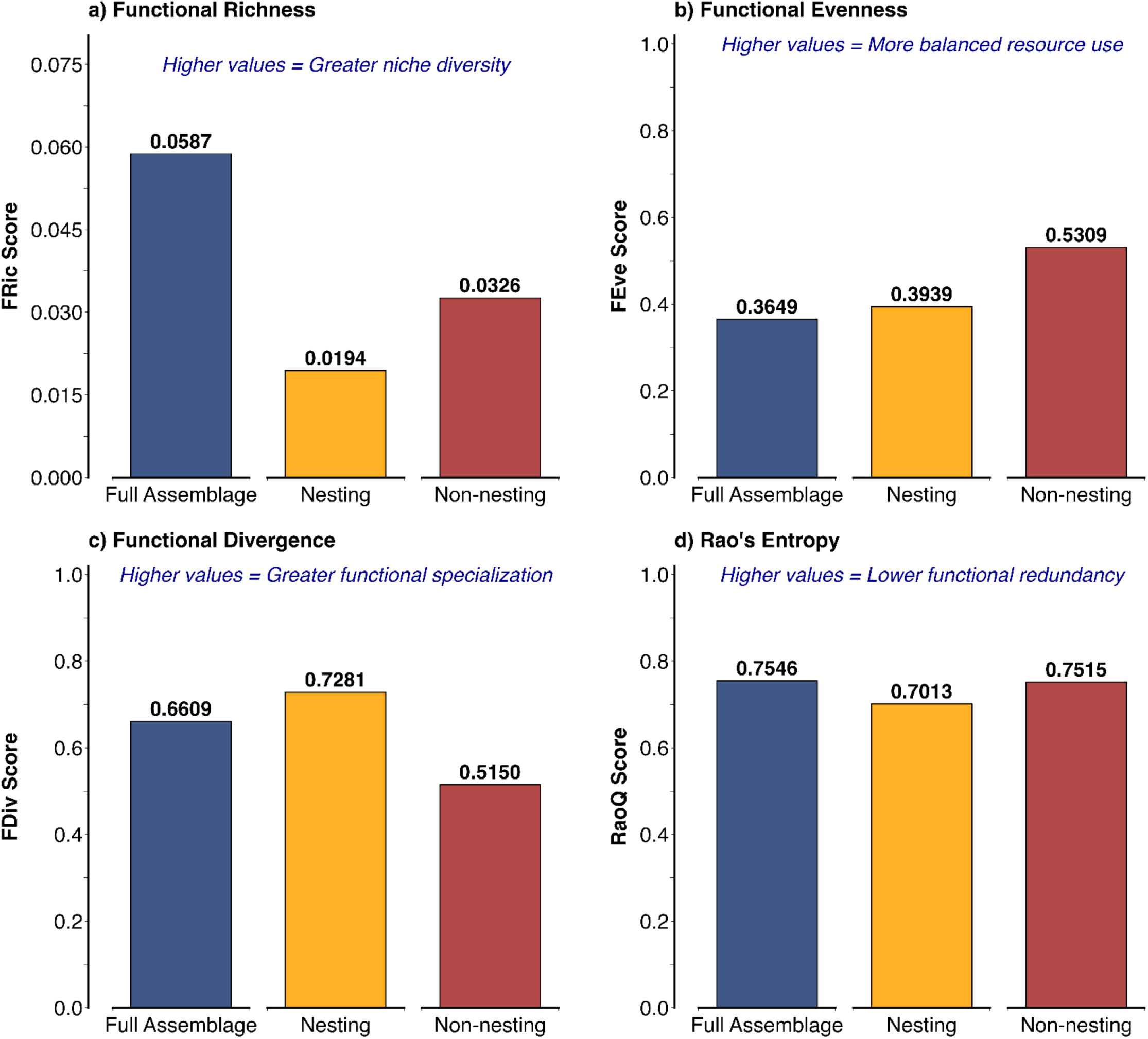
Functional diversity metrics for the full seabird assemblage and for nesting and non-nesting groups in the SGNRs EBSA. Panels show **(a)** functional richness (FRic), **(b)** functional evenness (FEve), **(c)** functional divergence (FDiv), and **(d)** Rao’s quadratic entropy (RaoQ).

Analysis of species-specific contributions to functional richness (FRic) and functional evenness (FEve) revealed strong asymmetry among taxa within both nesting and non-nesting seabird groups (Table 4). In the nesting group, FRic was overwhelmingly driven by Great Frigatebirds (contributing 48.9% of the total FRic value), followed by Christmas Shearwater (17.8%) and Masked Booby (8.9%). In contrast, among non-nesting species, the most significant contributions to FRic were from Chilean skua (40.7%), and Sooty Shearwater (22.2%). For FEve, Masked Booby (12.5%), Juan Fernandez Petrel (10.8%), and Phoenix Petrel and Great Frigatebird (9.3% and 9.1%, respectively) were the top contributors in the nesting assemblage, while FEve in the non-nesting group was most influenced by Wandering Albatross (16.5%), White-headed Petrel (11.3%), and White-chinned Petrel (9.8%).

To further explore functional differences between nesting and non-nesting groups, which were significant for FRic and FEve, we conducted an additional analysis of morphological traits and primary lifestyle categories. The selected traits –mass, wing length, tarsus length, beak length (culmen), and tail length– were chosen based on their substantial contribution to the functional diversity differences between groups, as identified in the trait contribution analysis (Table 3). These morphological traits are known to reflect functional roles in seabird ecology, such as foraging strategies and dispersal abilities, which align with the observed differences in functional diversity metrics. Non-nesting species had significantly higher values for mass, wing length, tarsus length, and beak length compared to nesting species, while no significant difference was observed for tail length (Figure 8a-e). In addition, the composition of primary lifestyle differed between groups, with differences in the proportion of aerial, aquatic, and generalist species (Figure 8f), further contributing to the functional divergence captured by the diversity indices. Together, these trait and lifestyle differences translated into a clear separation within the multivariate functional space, where several species pulled each group towards distinct regions that were not shared between them (Figure 9). For example, nesting species like the Masked Booby and Great Frigatebird extended towards the positive end of PC1 and PC2, highlighting traits associated with piscivory and aerial lifestyles. In contrast, among non-nesting species, the Southern Giant-Petrel defined the extremes along the positive end of PC1 and negative end of PC2, whereas Wilson’s Storm-Petrel defined the extremes along the negative PC1 and positive PC2, reflecting their distinctive generalist and aerial lifestyles, respectively.

**Figure 8.**
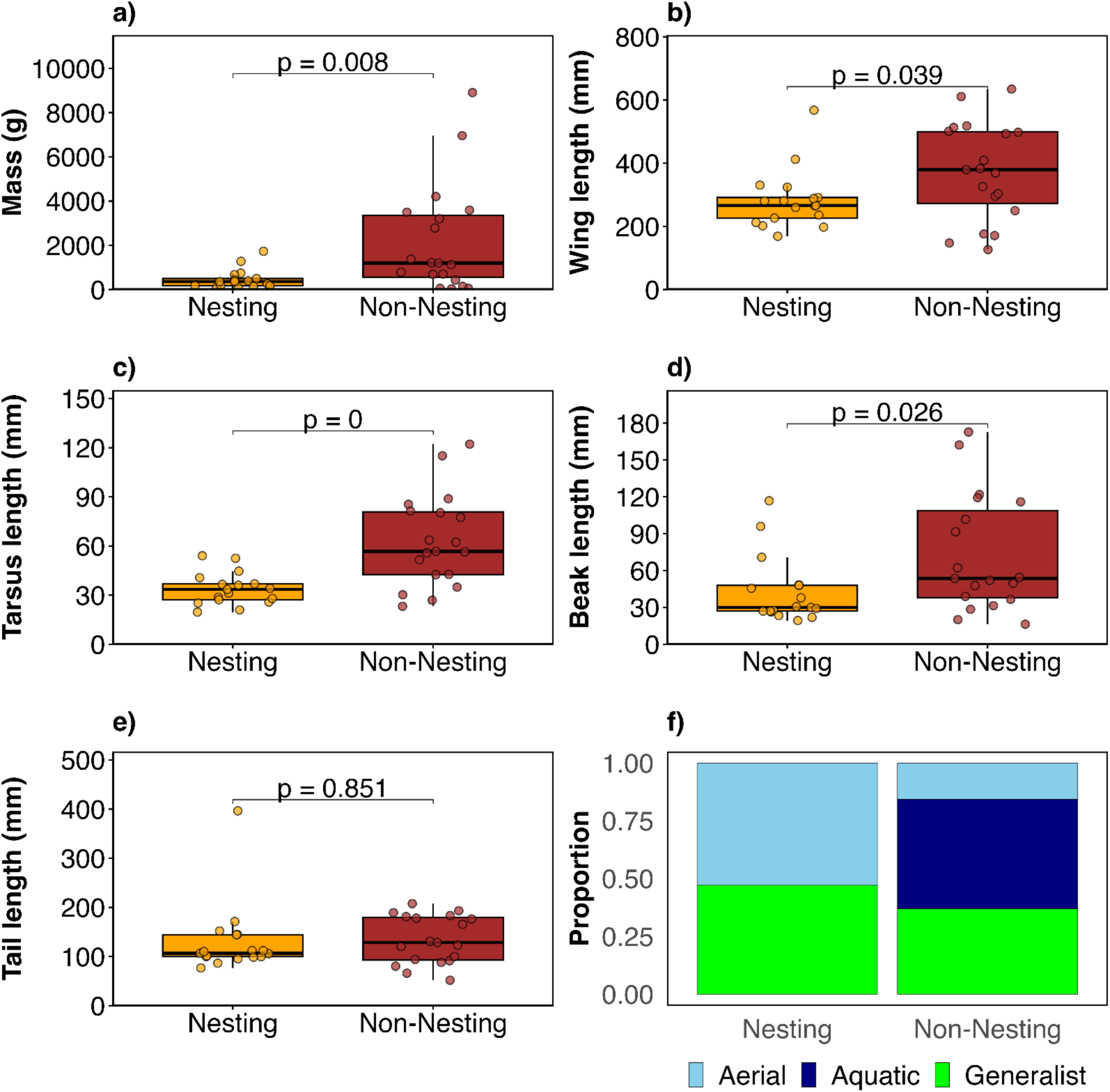
Comparison of key morphological traits and primary lifestyle proportions between Nesting and Non-Nesting seabird groups. Boxplots display the distribution of values for body mass (g), wing length (mm), tarsus length (mm), beak length (mm), and tail length (mm), with significant differences between groups indicated by p-values after a Student t-test with correction for unequal variances. The bar chart shows the proportional composition of primary lifestyles for each group, highlighting differences in functional strategies. Points correspond to the species and were jittered to avoid overlapping points.

**Figure 9.**
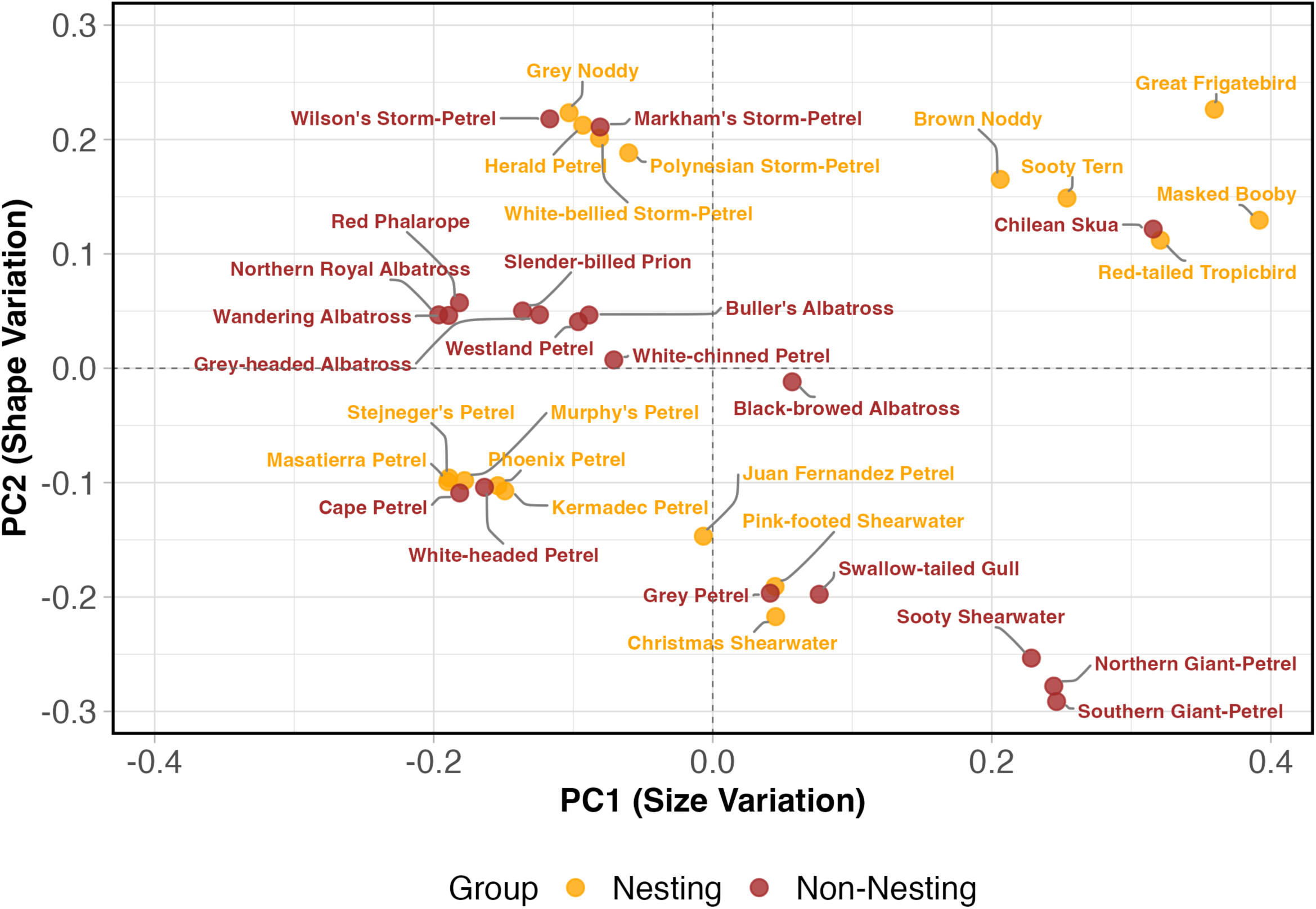
Functional trait space for nesting and non-nesting seabirds along the first two principal components. Each point represents a species, coloured by group: nesting (orange) and non-nesting (brown). The x-axis (PC1) mainly reflects variation in size-related traits, and the y-axis (PC2) reflects variation in shape-related traits. The common names are shown for each species.

**Table 3.** Results of trait-based comparisons between nesting and non-nesting seabird groups in the Salas & Gómez and Nazca Ridges EBSA. Significant differences (in bold) were found for PC1 associated to size variation, PC2 associated to shape variation, and Primary Lifestyle, which guided the selection of the specific morphological and lifestyle traits presented in Figure 5 and 6, as these traits contributed most to the functional diversity differences between groups.

| Trait | Statistic | p-Value | Test |
| --- | --- | --- | --- |
| PC1 Size variation | 93.00 | **0.0299** | Wilcoxon |
| PC2 Shape variation | 74.00 | **0.0048** | Wilcoxon |
| Primary Lifestyle | 11.99 | **0.0024** | Chi-square |
| Foraging strata: |  |  |  |
| on the ground | 136.00 | 0.0984 | Wilcoxon |
| around the surface | 170.00 | 0.7931 | Wilcoxon |
| below the surface | 178.50 | 0.5825 | Wilcoxon |
| Diet | 0.43 | 0.8065 | Chi-square |

**Table 4.** Relative contribution (%) of individual species to Functional Richness (FRic) and Functional Evenness (FEve) for nesting and non-nesting seabirds in the Salas & Gómez and Nazca Ridges.

| Functional Richness (Fric) |  |  |  | Functional Evenness (FEve) |  |  |  |
| --- | --- | --- | --- | --- | --- | --- | --- |
| Nesting species | % | Non-nesting species | % | Nesting species | % | Non-nesting species | % |
| Great Frigatebird | 48.9 | Chilean Skua | 40.7 | Masked Booby | 12.5 | Wandering Albatross | 16.5 |
| Christmas Shearwater | 17.8 | Sooty Shearwater | 22.2 | Juan Fernandez Petrel | 10.8 | White-headed Petrel | 11.3 |
| Masked Booby | 8.9 | Northern Royal Albatross | 3.4 | Phoenix Petrel | 9.3 | White-chinned Petrel | 9.8 |
| Red-tailed Tropicbird | 8.5 | Cape Petrel | 3.0 | Great Frigatebird | 9.1 | Grey Petrel | 7.8 |
| Pink-footed Shearwater | 4.7 | Wilson's Storm-Petrel | 1.7 | White-bellied Storm-Petrel | 8.1 | Buller's Albatross | 7.7 |
| Grey Noddy | 1.7 | Southern Giant-Petrel | 1.4 | Polynesian Storm-Petrel | 6.0 | Westland Petrel | 7.1 |
| Polynesian Storm-Petrel | 0.8 | Red Phalarope | 1.4 | Sooty Tern | 5.5 | Wilson's Storm-Petrel | 6.5 |
| Kermadec Petrel | 0.5 | Northern Giant-Petrel | 1.3 | Masatierra Petrel | 4.8 | Black-browed Albatross | 4.4 |
| Herald Petrel | 0.4 | Markham's Storm-Petrel | 1.1 | Stejneger's Petrel | 3.7 | Red Phalarope | 4.0 |
| Murphy's Petrel | 0.2 | Grey Petrel | 0.3 | Brown Noddy | 3.3 | Slender-billed Prion | 3.9 |
| Sooty Tern | 0.2 | Black-browed Albatross | 0.3 | Grey Noddy | 2.5 | Swallow-tailed Gull | 3.6 |
| Masatierra Petrel | 0.1 | Swallow-tailed Gull | 0.0 | Red-tailed Tropicbird | 1.3 | Cape Petrel | 3.0 |
| Stejneger's Petrel | 0.1 | Wandering Albatross | 0.0 | Murphy's Petrel | 1.1 | Northern Royal Albatross | 2.9 |
| Brown Noddy | 0.0 | Slender-billed Prion | 0.0 | Herald Petrel | 0.7 | Southern Giant-Petrel | 2.7 |
| Juan Fernandez Petrel | 0.0 | White-chinned Petrel | 0.0 | Christmas Shearwater | 0.3 | Northern Giant-Petrel | 1.9 |
| Phoenix Petrel | 0.0 | Westland Petrel | 0.0 | Pink-footed Shearwater | 0.2 | Sooty Shearwater | 1.4 |
| White-bellied Storm-Petrel | 0.0 | White-headed Petrel | 0.0 | Kermadec Petrel | 0.0 | Chilean Skua | 1.3 |
|  |  | Buller's Albatross | 0.0 |  |  | Grey-headed Albatross | 1.0 |
|  |  | Grey-headed Albatross | 0.0 |  |  | Markham's Storm-Petrel | 0.3 |

## Discussion

Our study provides the first comprehensive trait-based assessment of seabird functional diversity in the SGNRs, contributing to the growing number of studies examining seabird assemblage structure and function. While previous studies have examined functional traits in seabirds across various contexts –including extinction risk prediction based on biological traits^21^ and cross-ecosystem nutrient subsidies^47^– comprehensive multidimensional functional diversity assessments using the full suite of functional diversity indices remain limited for oceanic seabird assemblages, particularly in biodiversity hotspots beyond national jurisdiction like the SGNRs. By moving beyond species inventories and applying a multidimensional trait-based approach, we offer valuable insights into how functional diversity underpins ecosystem processes and resilience in this unique marine environment. These findings directly inform ongoing efforts to prioritise effective conservation strategies, not only highlighting species richness but also the importance of maintaining a full range of ecological functions.

While our study represents a significant advance in understanding seabird functional diversity, it also presents inherent limitations. First, our dataset is based primarily on at-sea observations, which, despite considerable survey effort, may under-represent particular elusive or research vessel-avoiding species^48^. The spatial coverage of our opportunistic at-sea transects likely missed parts of the SGNR region, and some key areas remain insufficiently surveyed; as a result, the absence of certain known breeders at sea may have led to localised underestimation of functional richness. Nevertheless, this dataset provides a high-resolution view of seabird assemblages at sea. Additionally, while our trait dataset was comprehensive in terms of morphology and foraging ecology, it did not include reproductive, energetic, or migratory traits, which could further refine understanding of functional roles and community vulnerability^16,49^. Also, dietary trait assignment relied on predominant prey types from global references and cannot address intraspecific or seasonal variation (*e.g.*, shifts between fish and squid), yet this standardised approach enables meaningful comparisons across species and regions and underscores the value of future studies incorporating more flexible indicators of dietary ecology.

Notwithstanding these limitations, our findings provide valuable perspectives on seabird functional diversity in the marine ecosystem of the SGNRs. By integrating at-sea abundance data with multidimensional trait analyses, we reveal a seabird assemblage characterized by low functional richness, moderate-to-low evenness, and high divergence. These patterns are significantly non-random and have important implications for ecosystem functioning and conservation^50^. The seabird assemblage displayed a compact yet heterogeneous trait structure, with most species (20 out of 36) clustering centrally, suggesting broadly shared ecological strategies, and a few occupying the margins of the trait space. Sixteen marginal species exhibited extreme combinations of traits related to diet, foraging behaviour, or body size, driving high functional divergence despite moderate overall richness. This result highlights the simultaneous presence of ecological redundancy and functionally distinctive species within the assemblage, supporting our first hypothesis that the SGNRs seabird assemblage is characterized by low functional richness and an uneven distribution of ecological roles arising from a constrained trait range and pronounced ecological structuring.

Our analyses demonstrate that the overall seabird assemblage exhibits extremely low functional richness (FRic), occupying a limited portion of the available trait space. Null model analyses confirm that both nesting and non-nesting groups have significantly lower functional richness than expected by chance, indicating strong ecological clustering and limited exploration of potential functional niches^50^. Functional evenness (FEve) is also low in nesting species and high in non-nesting species relative to null expectations, suggesting that nesting seabirds are more functionally clustered, while non-nesting seabirds are more evenly distributed across trait space.

Whether this constrained functional structure reflects severe environmental filtering alone, or is also shaped by historical processes such as the local extinction of breeding species that may once have occupied additional functional niches (*e.g*., benthic or deep-diving foragers), remains an open question. Although we lack paleontological or habitat-availability data to test these alternatives, disentangling contemporary filtering from historical legacies represents a promising direction for future research in this region.

Functional divergence (FDiv) was notably higher in nesting seabirds than in non-nesting ones reflecting the presence of specialized ecological roles among nesting taxa, although this difference did not reach statistical significance under the null model expectations. In contrast, non-nesting seabirds exhibit lower divergence, with species more concentrated toward the centre of trait space. These results support our second hypothesis, which predicted that functional diversity would differ between groups, with nesting seabirds showing higher functional divergence and non-nesting seabirds greater functional richness and evenness. However, Rao’s quadratic entropy (RaoQ), which integrates both abundance and trait differences, showed similarly high values for nesting and non-nesting groups. This suggests that, despite some differentiation, the overall functional diversity is constrained in this marine ecosystem.

Morphological traits (body mass, wing and beak length, tarsus length) and primary lifestyle (aerial, aquatic, generalist) are the main drivers of functional differentiation between groups, consistent with previous findings^50^. In this oceanic system, non-nesting species are generally larger and exhibit a broader range of lifestyles, likely contributing to their higher functional richness and evenness. In contrast, nesting species, while more functionally divergent, are restricted to a narrower trait space, possibly due to ecological constraints associated with breeding. During this period, adults are constrained by place-central foraging requirements, as they must return to the nest for incubate eggs and provision chicks, thereby facing constraints that may be exacerbated by intra– and interspecific competition for prey in the waters surrounding nesting colonies^11^.

The moderate Bhattacharyya Affinity (61%) between nesting and non-nesting groups indicates that, while there is substantial overlap in functional roles, each group also contains unique trait combinations not shared by the other. Although some degree of functional redundancy –the extent to which different species perform similar ecological functions, such that the loss of one may be compensated by others– exists among species occupying central trait space, the high proportion of vertex species and the high functional divergence suggest overall low redundancy, implying that the loss of certain species would likely translate into the loss of unique functional roles. However, low redundancy does not imply ecological equivalence, and preserving taxa with unique trait combinations remains essential, especially in the context of increasing human pressures in areas beyond national jurisdiction where seabird populations remain largely unprotected^51,52^.

The pronounced asymmetry in species’ contributions to functional richness (FRic) and evenness (FEve) observed between nesting and non-nesting seabird groups provides key insights into the ecological and evolutionary structuring of these assemblages. In nesting seabirds, FRic was overwhelmingly shaped by a small subset of taxa –most notably Great Frigatebird, followed by Christmas Shearwater and Masked Booby. These species exhibit distinctive trait combinations related to extreme aerial foraging abilities (*e.g.*, high aspect ratio wings in Great Frigatebird, plunge-diving in Masked Booby) or specific body size and life-history combinations (as in Christmas Shearwater). Their dominant influence on FRic suggests that evolutionary specialisations for nesting and foraging on isolated oceanic islands facilitate occupation of unique functional niches, expanding the range of functional space despite limited species richness. Conversely, among non-nesting seabirds, FRic was more evenly apportioned among species such as Chilean Skua and Sooty Shearwater, each characterised by their distinctive foraging ecologies and life-history strategies. This pattern likely reflects the diverse evolutionary origins and dispersal histories of non-nesting taxa, some of which migrate into the study region and thus enrich trait space from multiple phylogenetic lineages.

For FEve, a similar pattern emerged: in the nesting group, four species –Masked Booby, Juan Fernandez Petrel, Phoenix Petrel and Great Frigatebird– substantially increased evenness by occupying otherwise unrepresented regions of trait space. In non-nesting species, Wandering Albatross, White-headed Petrel and White-Chinned Petrel played the most significant roles, likely due to their unique intersections of large body size, broad foraging strata, and foraging plasticity. Such asymmetrical contributions shape the overall functional structure, indicating that specific lineages have a disproportionate effect on both the breadth and regularity of functional trait distributions.

Taken together, these findings underscore the fundamental role of a few functionally distinctive species in maintaining the boundaries and redundancy of ecological trait space. The evolutionary pathways that have led to divergence among these key taxa –particularly in adaptation to local breeding, foraging, and movement constraints– underline the importance of conserving not only species richness but the phylogenetic and functional distinctiveness that secures ecosystem functioning in oceanic systems.

### Ecological and Conservation Implications

The Salas y Gómez and Nazca Ridges seabird assemblage exhibits functional traits distinct from those of other major oceanic systems, underscoring its ecological singularity. Compared to the Humboldt Current System –where seabird assemblages show higher functional richness driven by upwelling-induced productivity and niche differentiation^6,30^– the SGNRs display pronounced functional clustering, with nesting species occupying a narrower trait space and non-nesting species showing higher evenness. This patterns also differs from the Canary Current Large Marine Ecosystem, where seabird diversity hotspots correlate with moderate productivity and high chlorophyll-*a* concentrations^53^. In the SGNRs, seabirds show relatively low functional redundancy but intermediate divergence, only partially resemble patterns observed in remote Polynesian islands. Whereas redundancy in Polynesian seabird communities has been interpreted as an adaptation to oligotrophic conditions^30^, the SGNRs’ low redundancy suggests that few species share similar functional roles. Although intermediate divergence still indicates some degree of niche differentiation^47^, the limited overlap among species may reduce the potential buffering of ecosystem processes against species loss.

Seabird diversity and biomass are both crucial for maintaining cross-ecosystem nutrient subsidies, with diversity especially important for sustaining offshore nutrient flows and ecosystem functioning globally^47^. At the SGNR, several seabird species that make a substantial contribution to functional richness illustrate how the loss of particular functional types could disrupt these processes. For instance, sympatric Gadfly petrels (*Pterodroma* spp.) occupy distinct isotopic niches, reflecting differences in spatial foraging distribution and trophic level^54^. In this context, Masked Booby forage mainly in the waters surrounding the islands, acting as a large plunge-divers that targets ephemeral fish aggregations near their breeding colonies^8^. Finally, Great Frigatebirds forage over broad oceanic areas and occasionally engage in aerial kleptoparasitism, which, although representing a minor component of its overall foraging strategy^55^, can still influence the foraging success and competitive interactions of other seabirds at sea^11^. Such niche partitioning implies that each species forages partially in different water masses and trophic levels, thereby conveying marine-derived nutrients from complementary at-sea areas back to the islands. Together, they function as a multi-species nutrient-transport network, integrating productivity across latitudinal and trophic gradients. Thus, the loss of any one species would be expected to selectively remove its specific nutrient pathway to the islands^56^. The positive effects of functional diversity on ecosystem processes highlight the need to protect not only species numbers but also the range of functional roles present in these communities^51^.

### Unique Vulnerabilities and Global Significance

The constrained functional diversity and low redundancy observed in SGNR seabird assemblages highlight a distinct vulnerability to environmental change and anthropogenic pressures^3^. The SGNRs’ seabirds face compounded threats due to the region’s jurisdictional complexity: 73% lies in areas beyond national jurisdiction, exposing it to unregulated fishing, plastic pollution, and seabed mining^2^. Unlike the well-studied Galápagos or Hawaiian archipelagos, where marine protected areas buffer localized threats, the SGNRs’ open-ocean foraging grounds lack coordinated protection, amplifying risks to wide-ranging species such as albatrosses and petrels^52^. The relatively low functional redundancy observed here –especially when contrasted with the higher redundancy reported for remote Polynesian islands and, more regionally, for seabird assemblages in the Humboldt Current System– suggest that the loss of seabird species would likely translate into the loss of unique functional roles, potentially compromising ecosystem processes such as cross-system nutrient subsidies^47^. Consistent with this, the intermediate functional divergence indicates that not all species are functionally equivalent, and the loss of certain unique species could disrupt key ecosystem functions^51^.

These findings underscore the need for conservation strategies that preserve not only species numbers but also the full spectrum of functional roles, particularly in regions of high endemism and ecological significance, where close to half of the species in many taxonomic groups –particularly seamount-associated fishes and benthic invertebrates– are endemic. Creating a marine protected area for the SGNRs under the BBNJ Treaty would promote coordinated management, strengthen ecosystem resilience, and help safeguard unique biodiversity in areas beyond national jurisdiction^2^.

## Author Contributions

GL-J conceived the study; PN directed the study, analysed the data and drafted several versions of the manuscript; PN and GL-J drafted the final version of the manuscript.

## Data availability

The data used in this study are available in Table 1, which contains data on abundance by species and their primary lifestyle, and Table S1, which contains information on biological traits.

## Acknowledgements

We would like to thank the Comité Oceanográfico Nacional (CONA) from Chile for funding several cruises under the CIMAR Islas Oceánicas Program and the Armada de Chile for permitting us to collect data on board their cruises. N. Luna, M. Portflitt-Toro, and J. Serratosa worked on board to collect, then maintain, and curate the data of our Seabird at Sea Monitoring Program database. J. B. Gusmao reviewed the first version of the manuscript. We appreciate the comments and suggestions from the reviewers, which contributed significantly to improving this study.

