## Supplementary material for "The Salas y Gomez and Nazca Ridges EBSA support a highly functional diversity of seabirds": Suplementary Material

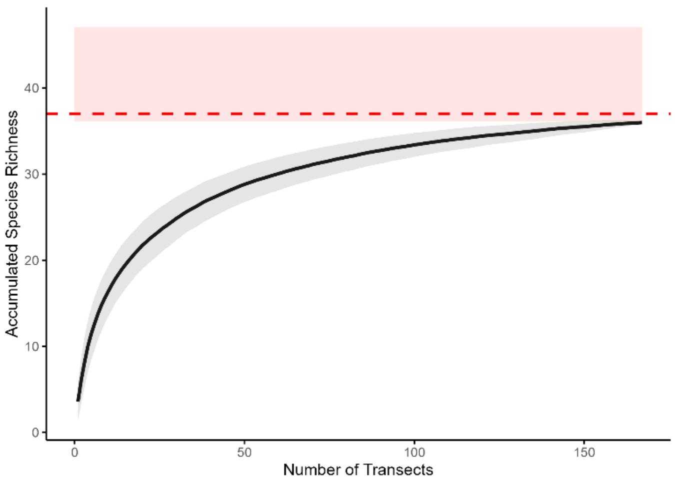


Supplementary Figure S1. Species accumulation curve for seabirds in the study EBSA area of Salas y Gómez and Nazca Ridge. The trend-smoothed black line was produced from 999 permutations using the random method. The shaded area represents the 95% confidence interval. The dashed red line indicates the estimated total species richness of 37 species, based on the Chao1 estimator, and the translucent red band corresponds to its 95% confidence interval (36–47 species).

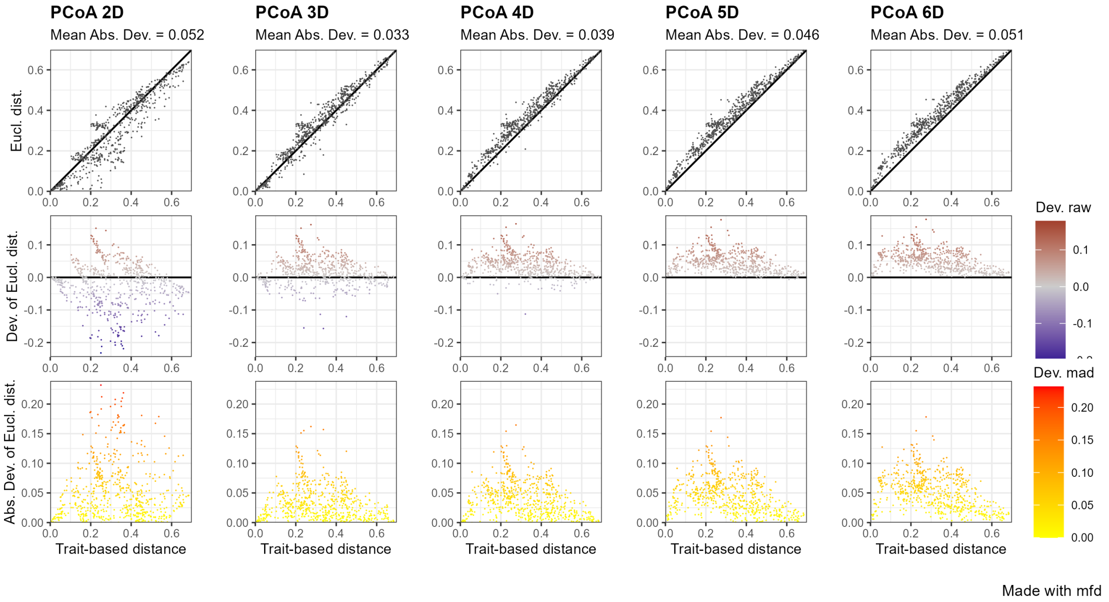


Supplementary Figure S2. Evaluation of the quality of selected functional spaces across different dimensionalities. The 3D space (PCoA_3D) exhibited the best quality with minimal deviation (MAD) in trait-based distances, effectively capturing the main ecological gradients and supporting its use for subsequent analyses. Gradient colors represent deviations in quality (yellow to red for decreasing quality) and trait distance deviations (dark blue to dark red).

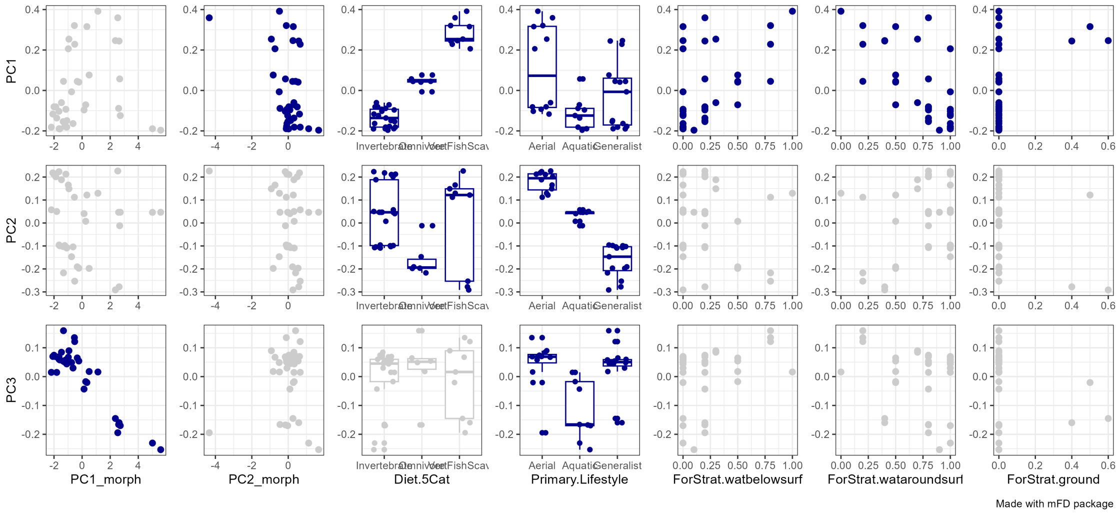


Supplementary Figure S3. Correlation between the functional traits of seabirds and PCoA axes. Plots in dark blue indicate significative correlations ($p < 0.05$).

**Table S1.** Seabirds recorded in the Salas y Gómez and Nazca Ridges. The status (nesting or non-nesting) and the traits used to calculate functional diversity are indicated. Foraging strata and diet were extracted from Wilman et al.[34], and morphology and lifestyle from Tobias et al.[33].

| Common name | Species | Status | Foraging Strata (%) Below Surface | Around Surface | On ground | Diet type |
|---|---|---|---|---|---|---|
| Brown Noddy | *Anous stolidus* | Nesting | 0 | 100 | 0 | VertFishScav |
| Pink-footed Shearwater | *Ardenna creatopus* | Nesting | 50 | 50 | 0 | Omnivore |
| Sooty Shearwater | *Ardenna griseus* | Non-nesting | 80 | 20 | 0 | VertFishScav |
| Swallow-tailed Gull | *Creagrus furcatus* | Non-nesting | 50 | 50 | 0 | Omnivore |
| Cape Petrel | *Daption capense* | Non-nesting | 0 | 100 | 0 | Invertebrate |
| Wandering Albatross | *Diomedea exulans* | Non-nesting | 0 | 100 | 0 | Invertebrate |
| Northern Royal Albatross | *Diomedea sanfordi* | Non-nesting | 10 | 90 | 0 | Invertebrate |
| Great Frigatebird | *Fregata minor* | Nesting | 20 | 80 | 0 | VertFishScav |
| White-bellied Storm-Petrel | *Fregetta grallaria* | Nesting | 20 | 80 | 0 | Invertebrate |
| Markham's Storm-Petrel | *Hydrobates markhami* | Non-nesting | 20 | 80 | 0 | Invertebrate |
| Southern Giant-Petrel | *Macronectes giganteus* | Non-nesting | 0 | 40 | 60 | VertFishScav |
| Northern Giant-Petrel | *Macronectes halli* | Non-nesting | 20 | 40 | 40 | VertFishScav |
| Polynesian Storm-Petrel | *Nesofregetta fuliginosa* | Nesting | 30 | 70 | 0 | Invertebrate |
| Wilson's Storm-Petrel | *Oceanites oceanicus* | Non-nesting | 0 | 100 | 0 | Invertebrate |
| Sooty Tern | *Onychoprion fuscatus* | Nesting | 30 | 70 | 0 | VertFishScav |
| Slender-billed Prion | *Pachyptila belcheri* | Non-nesting | 20 | 80 | 0 | Invertebrate |
| Red-tailed Tropicbird | *Phaethon rubricauda* | Nesting | 80 | 20 | 0 | VertFishScav |
| Red Phalarope | *Phalaropus fulicarius* | Non-nesting | 0 | 100 | 0 | Invertebrate |
| White-chinned Petrel | *Procellaria aequinoctialis* | Non-nesting | 50 | 50 | 0 | Invertebrate |
| Grey Petrel | *Procellaria cinerea* | Non-nesting | 50 | 50 | 0 | Omnivore |
| Westland Petrel | *Procellaria westlandica* | Non-nesting | 20 | 80 | 0 | Invertebrate |
| Grey Noddy | *Procelsterna albivitta* | Nesting | 0 | 100 | 0 | Invertebrate |
| Phoenix Petrel | *Pterodroma alba* | Nesting | 20 | 80 | 0 | Invertebrate |
| Masatierra Petrel | *Pterodroma defilippiana* | Nesting | 0 | 100 | 0 | Invertebrate |
| Juan Fernandez Petrel | *Pterodroma externa* | Nesting | 0 | 100 | 0 | Omnivore |
| Herald Petrel | *Pterodroma heraldica* | Nesting | 0 | 100 | 0 | Invertebrate |
| White-headed Petrel | *Pterodroma lessoni* | Non-nesting | 0 | 100 | 0 | Invertebrate |
| Stejneger's Petrel | *Pterodroma longirostris* | Nesting | 0 | 100 | 0 | Invertebrate |
| Kermadec Petrel | *Pterodroma neglecta* | Nesting | 20 | 80 | 0 | Invertebrate |
| Murphy's Petrel | *Pterodroma ultima* | Nesting | 0 | 100 | 0 | Invertebrate |
| Christmas Shearwater | *Puffinus nativitatis* | Nesting | 80 | 20 | 0 | Omnivore |
| Chilean Skua | *Stercorarius chilensis* | Non-nesting | 0 | 50 | 50 | VertFishScav |
| Masked Booby | *Sula dactylatra* | Nesting | 100 | 0 | 0 | VertFishScav |
| Buller's Albatross | *Thalassarche bulleri* | Non-nesting | 20 | 80 | 0 | Invertebrate |
| Grey-headed Albatross | *Thalassarche chrysostoma* | Non-nesting | 0 | 100 | 0 | Invertebrate |
| Black-browed Albatross | *Thalassarche melanophris* | Non-nesting | 20 | 80 | 0 | Omnivore |

**Table S1.** Continuation.

| Common name | Primary Lifestyle | Beak Length (mm) | Tarsus Length (mm) | Wing Length (mm) | Tail Length (mm) | Mass (g) |
|---|---|---|---|---|---|---|
| Brown Noddy | Aerial | 45.6 | 25.8 | 265.4 | 145.2 | 177.8 |
| Pink-footed Shearwater | Generalist | 48.4 | 52.5 | 324.0 | 112.2 | 744.0 |
| Sooty Shearwater | Generalist | 47.8 | 55.8 | 295.6 | 88.0 | 787.0 |
| Swallow-tailed Gull | Generalist | 62.2 | 51.5 | 409.0 | 183.5 | 687.0 |
| Cape Petrel | Generalist | 36.6 | 42.6 | 249.8 | 94.0 | 429.6 |
| Wandering Albatross | Aquatic | 162.3 | 115.1 | 634.5 | 181.3 | 6961.3 |
| Northern Royal Albatross | Aquatic | 172.8 | 122.2 | 610.8 | 176.5 | 8905.6 |
| Great Frigatebird | Aerial | 116.8 | 19.6 | 567.8 | 396.5 | 1274.7 |
| White-bellied Storm-Petrel | Aerial | 19.3 | 36.9 | 168.6 | 76.8 | 50.6 |
| Markham's Storm-Petrel | Aerial | 20.1 | 23.2 | 170.8 | 91.4 | 53.1 |
| Southern Giant-Petrel | Generalist | 91.6 | 85.4 | 493.6 | 178.2 | 3591.7 |
| Northern Giant-Petrel | Generalist | 101.7 | 88.9 | 498.0 | 165.8 | 4206.3 |
| Polynesian Storm-Petrel | Aerial | 21.9 | 44.6 | 197.6 | 99.8 | 70.0 |
| Wilson's Storm-Petrel | Aerial | 16.3 | 34.9 | 147.2 | 66.0 | 30.4 |
| Sooty Tern | Aerial | 47.9 | 21.0 | 280.7 | 152.1 | 185.7 |
| Slender-billed Prion | Aquatic | 28.5 | 30.2 | 175.8 | 80.4 | 145.0 |
| Red-tailed Tropicbird | Aerial | 70.8 | 28.7 | 330.5 | 99.8 | 671.9 |
| Red Phalarope | Aquatic | 31.4 | 26.9 | 126.3 | 51.8 | 55.6 |
| White-chinned Petrel | Aquatic | 54.4 | 62.3 | 382.6 | 120.4 | 1213.0 |
| Grey Petrel | Generalist | 49.7 | 56.7 | 325.8 | 99.8 | 1131.0 |
| Westland Petrel | Aquatic | 52.0 | 56.4 | 368.9 | 131.0 | 1199.0 |
| Grey Noddy | Aerial | 30.0 | 25.2 | 201.8 | 106.5 | 72.2 |
| Phoenix Petrel | Generalist | 26.3 | 31.2 | 266.2 | 101.2 | 259.0 |
| Masatierra Petrel | Generalist | 27.5 | 27.9 | 226.0 | 95.4 | 169.0 |
| Juan Fernandez Petrel | Generalist | 37.8 | 36.1 | 291.3 | 144.3 | 428.0 |
| Herald Petrel | Aerial | 27.0 | 34.1 | 281.0 | 110.0 | 394.0 |
| White-headed Petrel | Generalist | 38.9 | 42.5 | 303.6 | 123.2 | 698.0 |
| Stejneger's Petrel | Generalist | 23.4 | 27.2 | 212.2 | 105.8 | 143.0 |
| Kermadec Petrel | Generalist | 30.6 | 36.6 | 288.0 | 98.5 | 501.0 |
| Murphy's Petrel | Generalist | 27.0 | 33.5 | 260.0 | 112.0 | 360.0 |
| Christmas Shearwater | Generalist | 29.1 | 40.7 | 235.5 | 86.5 | 356.0 |
| Chilean Skua | Aerial | 53.6 | 63.6 | 379.6 | 128.5 | 1367.8 |
| Masked Booby | Aerial | 96.0 | 54.0 | 412.2 | 171.2 | 1732.1 |
| Buller's Albatross | Aquatic | 121.9 | 77.5 | 501.2 | 207.6 | 2781.5 |
| Grey-headed Albatross | Aquatic | 116.0 | 81.4 | 518.0 | 189.2 | 3499.0 |
| Black-browed Albatross | Aquatic | 119.3 | 80.3 | 513.2 | 193.2 | 3202.9 |
